# Mitochondrial genome microhomology-mediated editing by donor DNA delivery into mitochondria in human cells

**DOI:** 10.1101/2025.11.02.686110

**Authors:** Vadim V. Maximov, Nikita Shebanov, Natalia Nikitchina, Rachel Rapoport, Yehoshua Maor, Ivan Tarassov, Ophry Pines, Nina Entelis

**Author notes:** Correspondence should be sent to V.V.M. and I.T. The presented research was conducted in Jerusalem, Israel and Strasbourg, France. V.V.M is currently a Senior Scientist at MitoCareX Bio Ltd, Ness Ziona, Israel. MitoCareX Bio Ltd has not been involved in the presented research by any mean and this company is only the current affiliation of V.V.M. V.V.M. – Department of Molecular Biology, Phytor Ltd, 19 Ya’akov El’azar St., Jerusalem, Israel. I.T. – UMR7156 Molecular Genetics, Genomics, Microbiology, CNRS/University of Strasbourg, 4, Allée Konrad Roentgen Strasbourg 67000, France.

## Abstract

Mutations in the mitochondrial DNA (mtDNA) are associated with severe human diseases, lacking efficient therapies. Direct correction of mtDNA mutations may offer a cure for such diseases. We propose a novel strategy based on double-stranded DNA (dsDNA) oligonucleotide delivery into mitochondria and intrinsic microhomology-mediated end joining (MMEJ) for mtDNA editing. This strategy enables introduction of multiple predefined nucleotide changes in mtDNA. For this, the presence of MMEJ activity in the human mitochondrial lysates was confirmed. 49 bp DNA oligonucleotide duplexes, fused to an RNA hairpin previously identified as a mitochondrial import signal, were delivered into the mitochondria of cultured human cells. Delivery of these donor dsDNA molecules, homological to an *ND4* site of mtDNA and bearing designed nucleotide changes, led to a low but statistically significant introduction of the designed nucleotide changes into mtDNA. Donor dsDNA delivery combined with the CRISPR/mito-AsCas12a system also resulted in a statistically significant number of an expected concomitant change of five nucleotides distributed across a 16-nucleotide *ND4* site of the mitochondrial genome. The proposed strategy may become an efficient mtDNA editing tool suitable for the correction of near-homoplasmic mutations such as Leber’s Hereditary Optic Neuropathy (LHON)-associated mutations in the *ND4* gene of mtDNA.

## Introduction

Mutations in mitochondrial DNA (mtDNA) are associated with a spectrum of health conditions including cancer, aging, and Parkinson’s disease^1,2^, in addition to their roles in primary mitochondrial diseases^3,4^. Leber’s hereditary optic neuropathy (LHON), which causes central vision loss at a young age, is the most common optic neuropathy and an important example of a mitochondrial disease^5^. The most frequent mutation 11778G>A, which is associated with LHON, occurs in the mitochondrial *NADH dehydrogenase subunit 4* (*ND4*) gene^6,7^ (Figure 1A) which is a component of the mitochondrial respiratory chain^8^. This mutation leads to R340H substitution in the ND4 protein^6^ (Figure 1A). Functional studies of disease-associated mtDNA mutations, as well as the study of the mammalian mitochondrial genome in general, were impeded by the lack of efficient mammalian mitochondrial genome editing tools. Such tools could be also applied for gene therapies against diseases caused by mutations in the mitochondrial DNA.

**Figure 1.**
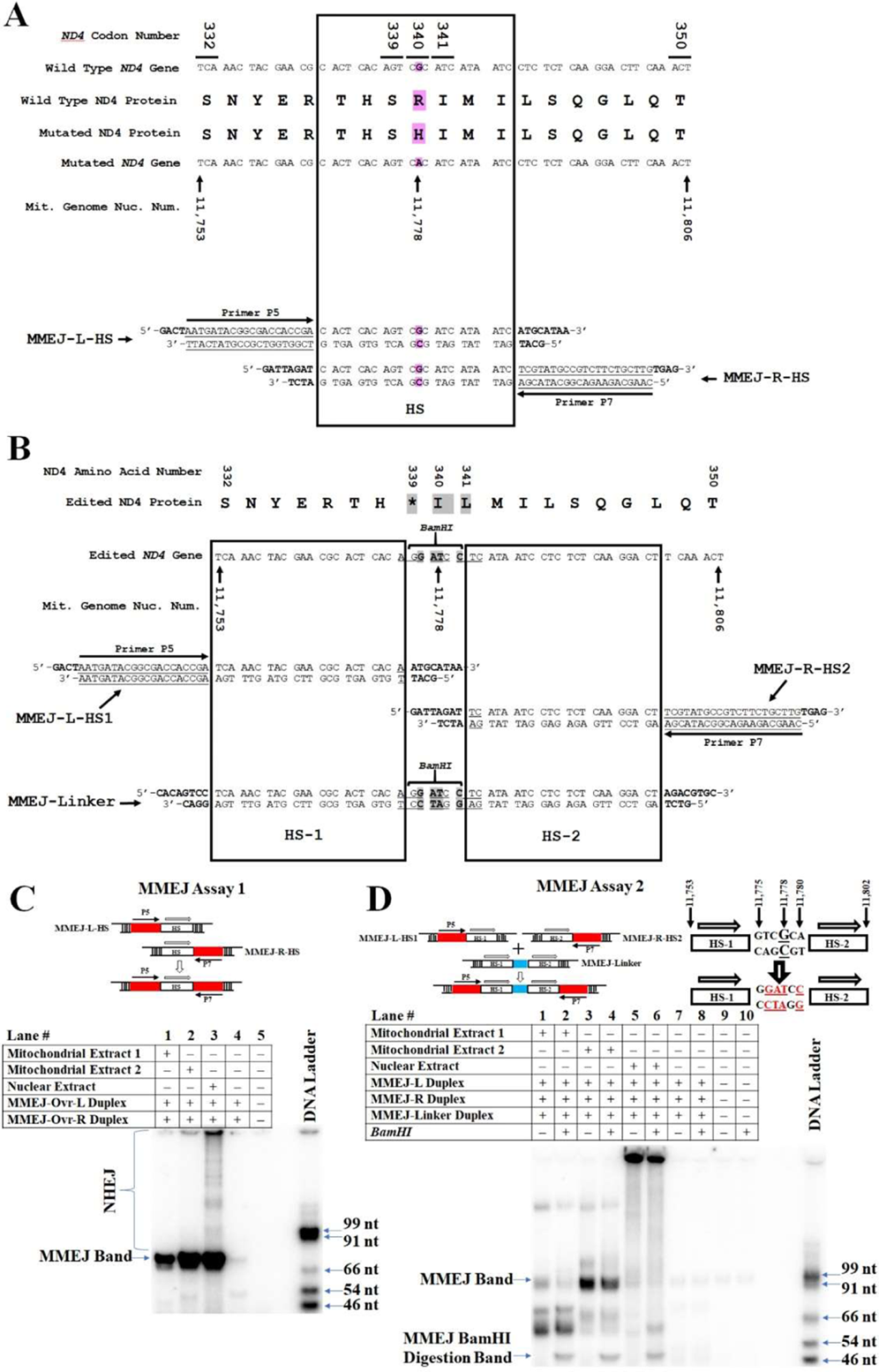
MMEJ Assay. (A) Sequences of the wild-type and LHON-associated mutant *ND4* gene alleles and ND4 proteins (upper panel) aligned to MMEJ-L-HS and MMEJ-R-HS duplexes (lower panel). Nucleotides and amino acid residues, which differ between wild-type and mutant alleles and proteins, are highlighted in purple. Sequence within the HS rectangle corresponds to the microhomology recombination site HS. (B) DNA duplexes applied for *BamHI* site insertion by MMJE (lower panel) and changes introduced in the *ND4* gene and ND4 protein during the mitochondrial genome editing by donor DNA delivery alone. The edited *ND4* gene nucleotides and corresponding amino acids are highlighted in bold gray. Sequences in the HS-1 and HS-2 rectangles correspond to the microhomology recombination sites HS-1 and HS-2. (C) End-joining activity in mitochondrial and nuclear extracts tested with DNA duplexes MMEJ-L-HS and MMEJ-R-HS (Supplementary Table 2) as schematically represented at the top. The MMEJ product was detected by radioactive PCR applying P5 and ^32^p-labeled P7 primers (highlighted in red) and separated by 10% urea PAGE. The HS sequence is shown on the panel (A). Bands above the MMEJ band correspond to the NHEJ products due to NHEJ activity from the nuclear extract (lane 3). (D) End joining activity tested in mitochondrial and nuclear extracts in presence of three DNA duplexes: MMEJ-L-HS1, MMEJ-Linker, and MMEJ-R-HS2, as schematically shown on the upper panel. The blue section in the middle of MMEJ-Linker represents the *BamHI* site. HS-1 and HS-2 sequences are shown on panel (B). Nucleotide G11778, which is commonly mutated in LHON, is underlined and enlarged. Nucleotides, which are different from those in the natural mitochondrial genome sequence and create the *BamHI* site, are in red and underlined. Autoradiography of amplicons, corresponding to MMEJ PCR product and its fragment (MMEJ *BamHI* Digestion Band) and separated by 10% urea PAGE, is shown. DNA Ladder represents a mix of ^32^p-labelled oligonucleotides, their size is indicated at the right.

Cytosine and adenosine deaminases-based tools have enabled the introduction of point transition nucleotide substitutions in mammalian mtDNA. Mitochondrially targeted rat APOBEC1, which can act as an RNA and DNA cytosine deaminase, was initially applied to introduce random cytidine to thymidine transition mutations in the Drosophila mitochondrial genome^9^. The first prominent success in mammalian mitochondrial genome editing was achieved by applying the double-stranded DNA deaminase toxin A (DddA)-derived cytosine base editor (CBE) (DdCBE)^10–19^. Mitochondrial adenosine base editors (ABEs) for efficient A-to-G editing, which are known as TALE-linked deaminases (TALEDs), have been also developed. TALEDs are fusions of TALE domains with a modified *E. coli* deoxyadenosine deaminase TadA—TadA8e^18,20,21^. Despite these impressive achievements, deaminase-based techniques have a fundamental limitation—they can introduce only point transition nucleotide changes (purine-to-purine or pyrimidine-to-pyrimidine).

All other modifications of the mammalian mitochondrial genome—transversion nucleotide changes (purine-to-pyrimidine or pyrimidine-to-purine), deletions, insertions, inversions, etc.— remain a challenge and require development of novel mitochondrial genome editing strategies. Mammalian mitochondria lack the classical non-homologous end joining (NHEJ)^22^, and linearized mtDNA is typically degraded by components of the replication machinery^23,24^. Remarkably, microhomology-mediated end joining (MMEJ), as well as homologous recombination activities, are detectable in mammalian mitochondrial extracts^22,25^. These activities could be potentially exploited for genome editing in mammalian mitochondria upon donor DNA delivery.

Delivery of linear DNA into isolated mammalian, including human, mitochondria has been reported^26,27^. However, DNA delivery into mitochondria of cultured mammalian cells has been a challenge. Amongst possible approaches are: (1) mitochondrial targeting peptides^28^, and (2) RNA mitochondrial import signals (RMISs)^29,30^. RMISs were discovered using SELEX^31^ and represent hairpin RNA structures that serve as a signal to target small non-coding RNA molecules in human mitochondria^32,33^. Chimeric RNA/DNA molecules, which contain RMIS and DNA sequences, can be commercially synthesized. RMIS, which is based on the D-arm of yeast 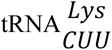, has been reported to successfully deliver not only RNA, but also short (20 nt) single-stranded DNA (ssDNA) oligonucleotides^29,30^. Nonetheless, it was unclear whether this RMIS can be applied for delivery of longer DNA duplexes. Here we report, for the first time, delivery of 49 nt DNA duplexes into mitochondria of cultured human cells, applying RMIS. Moreover, we demonstrate the possibility of human mitochondrial genome editing due to delivery of ss- and double-stranded DNA (dsDNA) oligonucleotides alone or in combination with CRISPR/mito-AsCas12a into mitochondria of human cells. Editing was carried out in the clinically relevant site of the mitochondrial *ND4* gene encompassing nucleotide G11778, whose mutations are associated with the LHON disease (Figure 1A).

## Results

### Microhomology-Mediated End Joining (MMEJ) in human mitochondrial extracts

To confirm the presence of MMEJ activity in human mitochondria^22^, we tested DNA end-joining activity in human mitochondrial and nuclear extracts. For this, we synthesized two oligonucleotide duplexes (Figure 1A, Table S1), bearing the same 22nt microhomology arm indicated as “HS” in Figures 1A and 1C. These duplexes were incubated with mitochondrial and nuclear extracts from HEK 293T cells. Recombination between microhomology arms of these sequences, resulting in joining their ends, indicates MMEJ activity. The recombination product is detectable by PCR (Figure 1C, MMEJ band) with primers P5 and P7 (Table S2). We detected MMEJ but not NHEJ activity, which is expected to give longer amplicons, in human mitochondrial extracts (Figure 1C, lanes 1 and 2), while both NHEJ and MMEJ activities could be detected in the human nuclear extract (Figure 1C, lane 3). This suggests that the detected mitochondrial MMEJ activity is not a result of nuclear contamination. The microhomology arm was purposely designed to be identical to the wild-type human mitochondrial genomic site, which is affected in LHON by the most common LHON mutation 11778G>A^6,7^ (Figure 1A), to test whether MMEJ recombination can occur at this site.

To further validate MMEJ activity in the mitochondrial lysate, we applied another test system consisting of three DNA fragments (Figure 1B, 1D and Table S1). Two DNA duplexes, MMEJ-L-HS1 and MMEJ-R-HS2, harbored different sequences HS-1 and HS-2 respectively. The third one, MMEJ-Linker, contained both HS-1 and HS-2 microhomology arms separated by *BamHI* site. Therefore, if two MMEJ-mediated recombination events successfully occur, this will produce a DNA fragment bearing the *BamHI* recognition site “GGATCC”, which replaces the wild-type site “GTC**<u>G</u>**CA” in the mitochondrial genome between HS-1 and HS-2 (Figures 1B, 1D). The resulting recombinant DNA was detected by PCR, and *BamHI* cleavage of the amplicon generated DNA fragments of the expected sizes (Figure 1D). These results suggest that the intrinsic mitochondrial MMEJ activity can be utilized for introduction of novel DNA sequences into the human mitochondrial genome.

### Delivery of dsDNA oligonucleotides into mitochondria in human cells

To validate the utility of mitochondrial MMEJ activity for mitochondrial genome editing in the living human cells, we next needed to target the DNA molecules into mitochondria. Previously, RNA mitochondrial import signals (RMIS) have been successfully used for delivery of artificial RNA and short (20 nt) ssDNA molecules into human mitochondria^29–31,34,35^. Here, we aimed to test whether an RNA hairpin, corresponding to the D-arm of yeast 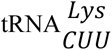, can be used to deliver longer (49 nt) dsDNA molecules, which are suitable for human mitochondrial genome editing by MMEJ. We designed chimeric oligonucleotides, which are referred to as RMIS-Dir and RMIS-Rev (Figure 2A). The 5’-RNA parts of these oligonucleotides are composed of 16 ribonucleotides, corresponding to the D-arm of yeast 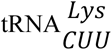 (in blue) where two dihydrouridine residues are substituted by uridine residues. The 3’-parts of the chimeric oligonucleotides represent 49 nt DNA sequence, corresponding to MMEJ-Linker (Figure 2A). These DNA sequences contain homologous arms, HS-1 and HS-2 (Figure 2A), which are required for recombination and integration in the human mitochondrial *ND4* gene. The five-nucleotide sequence at the center, together with the last nucleotide of the left flanking sequence, creates the six-nucleotide *BamHI* restriction site, which should replace the most common “LHON” mutation site in the mitochondrial genome. Thus, these oligonucleotides are designed to test whether the *ND4* region of human mitochondrial genome can be edited upon their delivery into human mitochondria. The expected nucleotide substitutions would lead to the changes in the ND4 protein: Ser339Stop, Arg340Ile, and Ile341Leu (Figure 1B). Therefore, these changes in the mitochondrial genome are expected to yield a truncated *ND4* protein and impair mitochondrial respiratory function.

**Figure 2.**
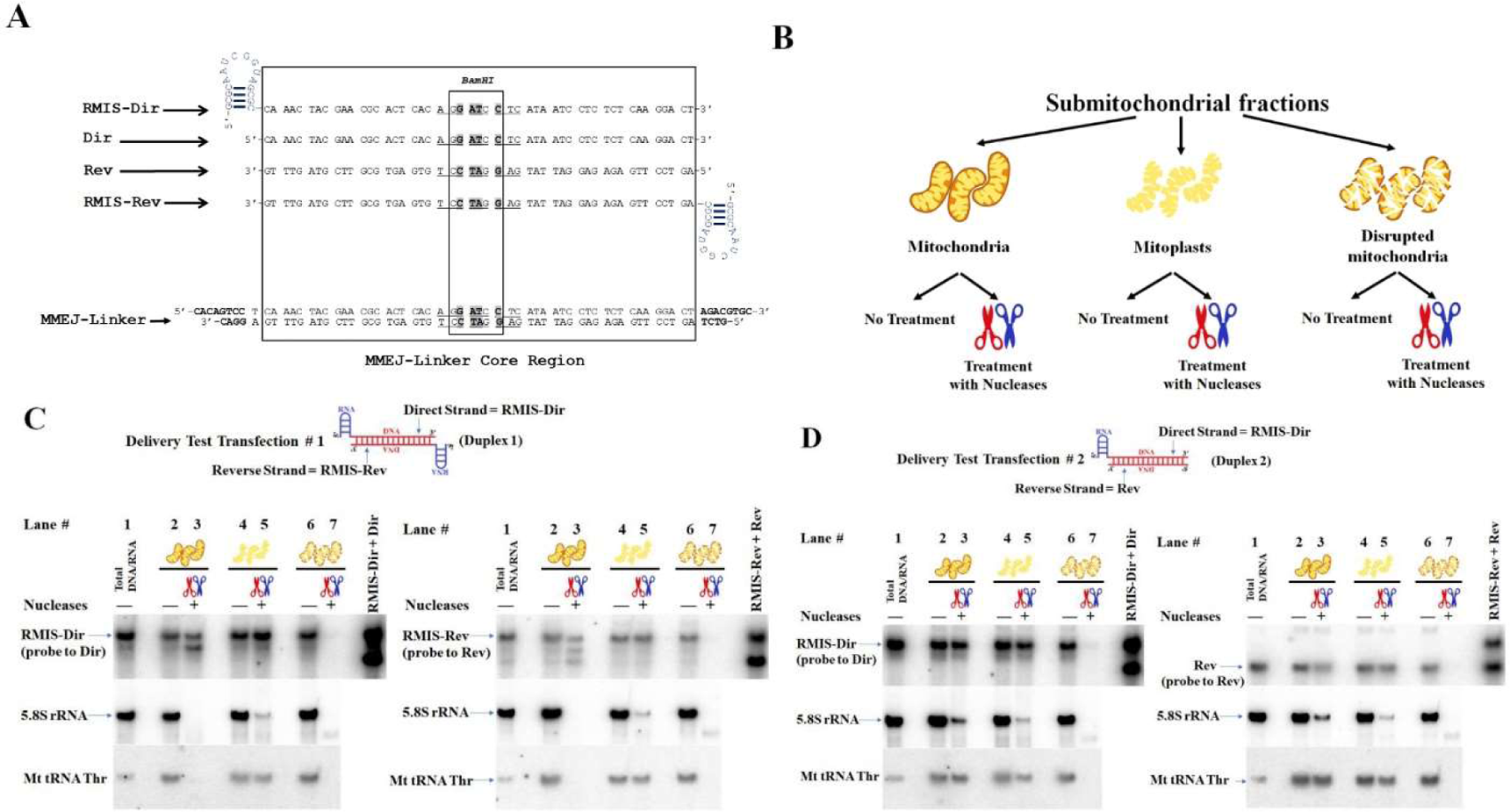
Evaluation of mitochondrial import of dsDNA fragments. (A) Chimeric RNA-DNA oligonucleotides RMIS-Dir, RMIS-Rev, as well as Dir and Rev, designed for mitochondrial delivery and editing experiments are shown in alignment to the MMEJ-Linker duplex. Nucleotides which differ from the wild-type *ND4* sequence are highlighted in gray. The *BamHI* site is shown by the small rectangle. Nucleotides, which differ from the wild-type sequence, are highlighted in bold. The blue hairpin structures represent RMIS. (B) Schematic representation of submitochondrial fractionation (modified from ^34^). (C, D) *In vivo* import Assays. Mitochondria were purified from the HEK 293T cells transfected with Duplex 1 RMIS-Dir:RMIS-Rev (C) or with Duplex 2 RMIS-Dir:Rev (D). Total nucleic acids (RNA + DNA) were purified and separated by 10% urea-PAGE for Northern blot hybridization. Two representative blots are shown for each experiment (left and right panels), probed for oligonucleotides Dir or Rev (upper left and right panels, correspondingly) as indicated on the left of each panel. To detect possible cytosolic contamination, the probe for 5.8S ribosomal RNA was used (middle panels); the mitochondrial tRNA^Thr^ signal (lower panel) served as a loading control and as an indicator of mitochondrial and mitoplast integrity. Mitochondria, mitoplasts and lysed mitochondria are schematically indicated above the panels, as in (B). Markers of size and hybridization controls, oligonucleotides RMIS-Dir and Dir, were loaded on the right extremity of each gel, as indicated on the top.

First, we asked whether dsDNA oligonucleotides with the RMIS on both strands (RMIS-Dir:RMIS-Rev, Figure 2C) or on only one strand (RMIS-Dir:Rev, Figure 2D) can be targeted into mitochondria in HEK 293T cells. For this, we modified the protocol previously developed to assess the mitochondrial import of small non-coding RNAs^31,32,36^. We transfected HEK 293T cells with the chimeric molecules and analyzed nucleic acids isolated from purified mitochondria, mitoplasts (obtained by swelling of mitochondria), and mitochondria lysed by detergent (Figure 2 B). To eliminate cytosolic nucleic acids, half of each fraction was treated with a mix of nucleases to ensure complete degradation of DNA duplexes outside the mitochondria (see Materials and Methods). Northern blot hybridization with probes to DNA sequences used for cell transfection (Dir and Rev), mitochondrial tRNA^Thr^ and cytosolic 5.8S rRNA demonstrated that nucleic acids located within the mitochondrial matrix were protected from the nuclease cleavage (Figure 2C, 2D, probe to mt tRNA^Thr^) while being sensitive to the nuclease treatment upon the mitochondrial membrane disruption. The 5.8S rRNA, localized in cytosolic ribosomes, was mostly sensitive to the nuclease treatment (Figure 2C, 2D). In one experiment, mitochondrial tRNA^Thr^ was degraded in mitochondria but not in mitoplasts (Figure 2C lanes 3 and 5), suggesting that mitochondria were accidently disrupted during purification. The chimeric duplexes (RMIS-Dir:RMIS-Rev and RMIS-Dir:Rev) were partially protected from the nuclease treatment in the mitoplast fractions (Figure 2C, 2D) and completely degraded by the nucleases upon lysis of mitochondria (Figure 2C, 2D).

Noteworthily, in cells transfected with Duplex 1 (Figure 2C), both strands of the duplex, RMIS-Rev and RMIS-Dir, were detected in the mitoplasts upon the nuclease treatment which indicate their delivery into the mitochondrial matrix. In cells transfected with Duplex 2 (Figure 2D, left panel), we clearly demonstrated the mitochondrial import of the chimeric RNA-DNA strand RMIS-Dir. Remarkably, the other strand of the duplex, Rev, not harboring the RMIS signal, was also protected from the nuclease treatment in the mitochondria and mitoplast fractions (Figure 2D, right panel), indicating delivery of this strand into mitochondrial matrix. These data demonstrate that a dsDNA duplex, with only one strand fused to an RNA hairpin structure, can penetrate mitochondrial membranes, thus expanding our knowledge of the nucleic acids mitochondrial import and its possible therapeutic applications.

### Mitochondrial genome editing in human cells by donor DNA delivery

Our next objective was to test whether mitochondrial import of donor DNA fragments can induce a change in human mtDNA sequence. For this, HEK 293T cells were transfected with a mix of ssDNA oligonucleotides RMIS-Dir and RMIS-Rev or with various pre-formed duplexes (Figure 3A) capable of being targeted into mitochondria. Six days post-transfection, mitochondria were isolated, and the targeted mtDNA region (an *ND4* gene site) was PCR-amplified and subjected to deep sequencing. Editing efficiency was defined as the number of reads containing the edited mitochondrial sequence per one million reads (Counts Per Million = CPM) that unambiguously aligned to the mitochondrial sequence (MT:11710-11855). Since multiple mitochondrial DNA insertions known as Nuclear mitochondrial Sequences (NumtS) are found in the human nuclear genome^37^, we applied the BLAT tool of the UCSC Genome Browser to search for sequences, which are similar to the sequenced amplicon, in human nuclear genome^38^. The most similar human nuclear genomic sequences (Table S3) were then aligned to the corresponding human mitochondrial genomic sequence (Figure S1A, see also Materials and Methods section for details). Ultimately, only those reads that clearly matched the mitochondrial but not the nuclear genome, were analyzed and used for the editing efficiency estimation (Figure 3B).

**Figure 3.**
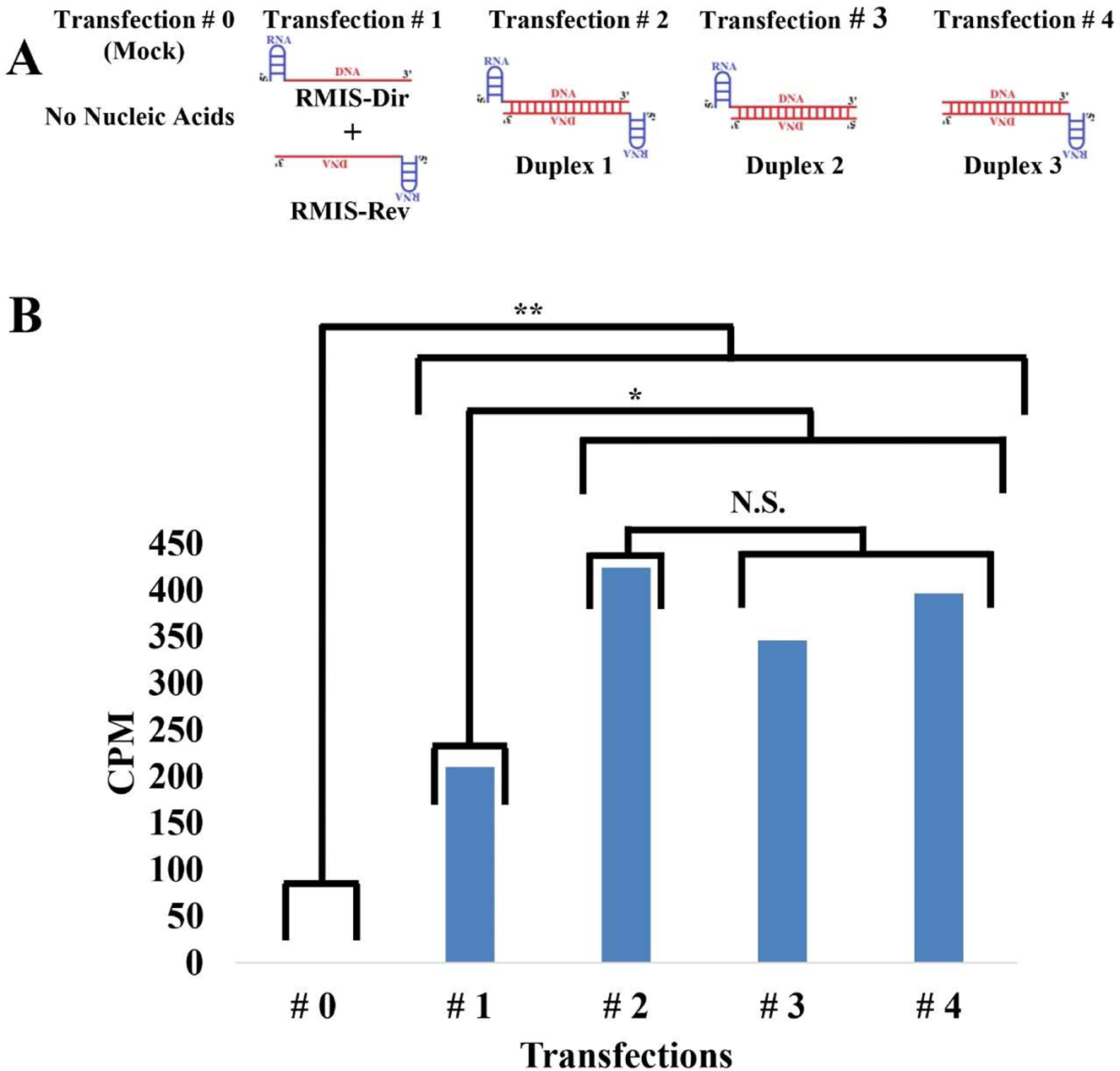
mtDNA editing by mitochondrially imported donor DNA in 293T cells. (A) Schematic representation of ss-oligonucleotides and duplexes used in the corresponding transfections of HEK 293T cells. Duplex 1 – RMIS-Dir:RMIS-Rev; Duplex 2 – RMIS-Dir:Rev; Duplex 3 – Dir:RMIS-Rev. (B) Mitochondrial genome editing efficiency in counts of reads with the edited mitochondrial sequence per one million (Counts Per Million = CPM) of reads unambiguously aligned to the mitochondrial sequence (MT:11713-11853) for each transfection is shown on the graph. Statistical significance was assessed by the two-sided permutation test with the plus-one correction and adjustment via Benjamini-Hochberg False Discovery Rate. N.S. – no statistical significance (p.adjust > 0.3). * – moderate statistical significance (p.adjust < 0.05; # 1 vs # 2 – p.adjust = 0.0036; # 1 vs # 3 – p.adjust = 0.043; # 1 vs # 4 – p.adjust = 0.0069). ** – strong statistical significance (p.adjust < 0.001).

We detected a low but statistically significant amount of the expected four-nucleotide change (see Figure 1C) only in samples from transfected cells (Figure 3B), indicating on a successful microhomology-mediated editing (further referred to as MME) of the targeted gene. Notably, RMIS-bearing DNA duplexes were significantly more efficient for human mitochondrial genome editing than the combination of single-stranded RMIS-Dir and RMIS-Rev oligonucleotides (see Discussion section). Thus, we succeeded to introduce the predesigned four-nucleotide change in the human mitochondrial genome in cultured human cells. It is important to mention that a four-nucleotide modification of the mitochondrial genome is currently not achievable by using other techniques, such as deaminase-based mitochondrial genome editing methods.

Next, trying to improve the efficiency of mitochondrial genome editing after donor DNA delivery, we decided to test whether mitochondrial DNA cleavage at the editing site can improve the MME efficiency. We used a recently constructed cell line, T-REx-293-Su9-AsCas12a, carrying a tetracycline-inducible gene of the mitochondria-targeted AsCas12a effector nuclease^39^. For editing, we selected a site within the mitochondrial *ND4* gene bearing the protospacer adjacent motif (PAM) TTTN required for recognition by the mito-AsCas12a effector nuclease (Figure 4A). For this experiment, new donor chimeric RNA/DNA oligonucleotides (RMIS-Dir-New and RMIS-Rev-New) were designed, each containing two different Homology Sequences regions (new HS-1 and new HS-2) flanking the site of specific cleavage by mito-AsCas12a/crRNA and the editing site (Figure 4A). A five-nucleotide change, which we expected to introduce by MME, is located in the region corresponding to crRNA, therefore, the donor DNA oligonucleotides and the edited mtDNA sequence should not be recognized and cleaved by the mito-AsCas12a/crRNA complex. Moreover, all the nucleotide changes were introduced in the third position of the codon, leading to synonymous codons and thus not inducing any alteration in the ND4 amino acid sequence.

**Figure 4.**
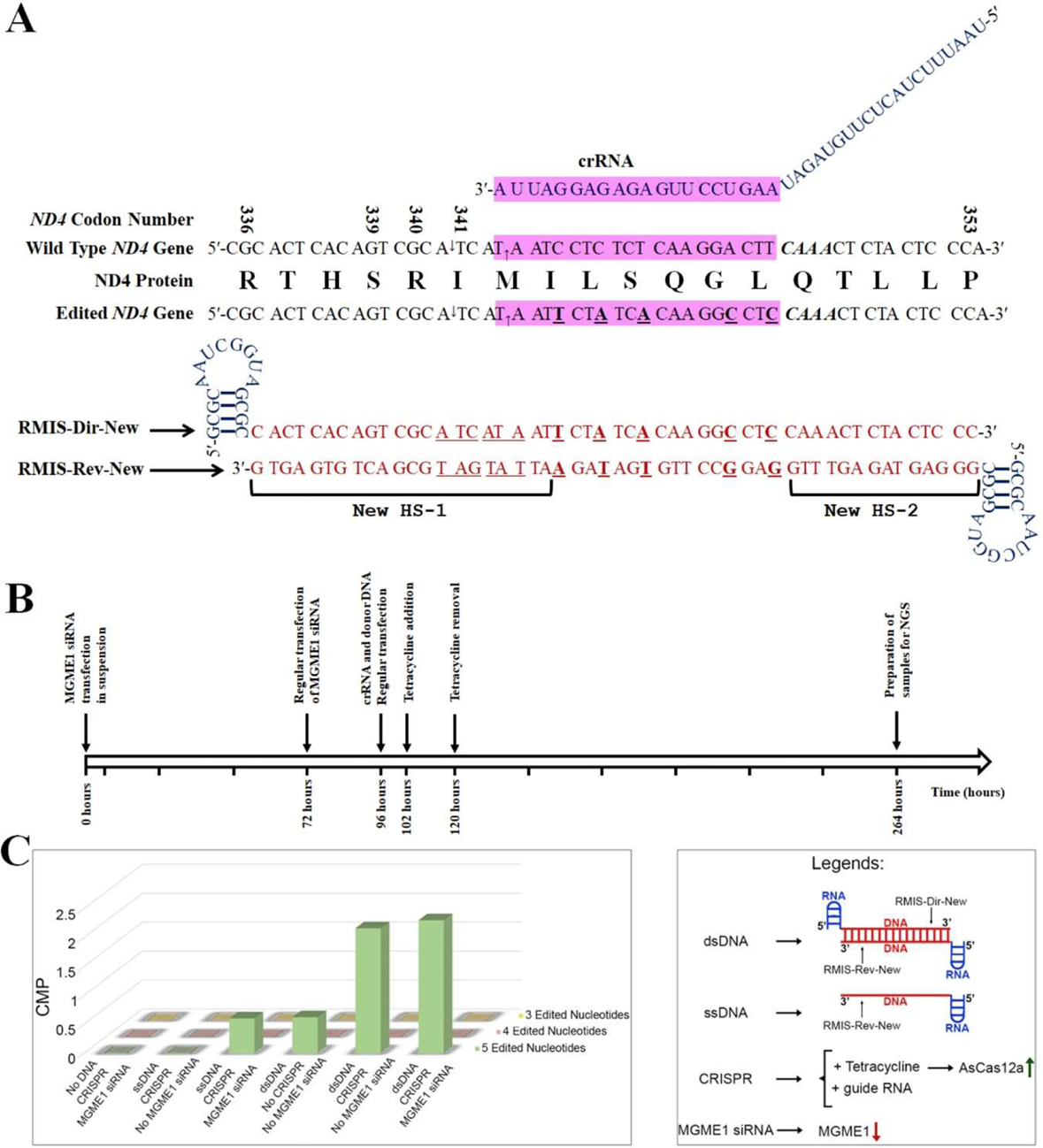
mtDNA editing by donor DNA delivery in mitochondria of T-REx-293-Su9-AsCas12a cells. (A) crRNA, wild-type and edited *ND4* alleles, and their translated protein sequences, RMIS-Dir-New and RMIS-Rev-New are shown on the picture. New HS-1 and New HS-2 indicate the microhomology arms. The guide part of crRNA and corresponding sites of wild-type and edited *ND4* genes are highlighted in purple. (B) Schematic representation of a timeline of cells transfections and treatments. (C) Left panel: means of counts of reads with five-, four-, or three edited nucleotides per one million (Counts Per Million = CPM) of reads unambiguously aligned to the mitochondrial sequence (MT:11643-11870) for each transfection, are shown on the 3D graph. Each transfection was conducted in biological triplicate. Right panel: schematic explanation of the transfections and treatments.

T-REx-293-Su9-AsCas12a cells were transfected with donor ssDNA or dsDNA together with the crRNA, in combination with induction of the mito-AsCas12a with tetracycline (Figure 4B, 4C). To prevent degradation of cleaved mtDNA molecules, some samples were also treated with an siRNA targeting *mitochondrial genome maintenance exonuclease 1* (*MGME1*), as described previously^39^. MGME1 plays a key role in the degradation of linearized mtDNA, thus, its downregulation was expected to stabilize the ends of cleaved mitochondrial DNA molecules^23^ and facilitate their recombination and ligation.

After cultivation of transfected cells, mtDNA was isolated and the region of interest was PCR-amplified with the introduction of unique molecular identifiers (UMIs)^40^. After the library preparation and deep sequencing, the editing efficiency was determined for different transfection conditions, including control experiment without donor DNA (Figure 4C). Differences between the mitochondrial sequence and similar NumtS outside the recombination region and the primers annealing sites (Figure S1B, Table S4) allow unambiguous alignment of reads to the mitochondrial sequence. Unexpectedly, we obtained extremely low levels of editing. To ensure the significance of the data, we compared the numbers of reads containing the expected five-nucleotide change and those with four- or three-nucleotide changes (Figure 1C and Figure S2). No four- or three-nucleotide changes were detected, indicating that the directed change of five nucleotides was not a random event. Moreover, no editing events were found in control cells without donor DNA delivery, suggesting that the editing requires donor DNA. The five-nucleotide change was also not found in cells transfected with ss-donor DNA without *MGME1* downregulation, which can indicate on the ssDNA degradation in mitochondria by MGME1.

The most significant level of mtDNA editing was obtained in cells transfected with ds-donor DNA and crRNA, both with and without *MGME1* silencing. Thus, as in the previous experiment, RMIS-bearing DNA duplexes were more efficient for human mitochondrial genome editing than the single-stranded chimeric oligonucleotides. Extremely low efficiency of editing in the last experiment may be explained by specific features of the T-REx-293-Su9-AsCas12a cell line, characterized by reduced respiration level^39^, which can influence the efficiency of cell transfection and the entire mitochondrial metabolism, including activity of enzymes involved in the microhomology mediated recombination.

Taken together, our results indicate that human mtDNA sequence can be modified by donor dsDNA delivery into mitochondria. This proof-of-concept study opens further possibilities to improve the efficiency of mtDNA editing and develop applications of this novel mtDNA editing approach.

## Discussion

### Challenge of mtDNA editing

The objective of the study was to apply the mitochondrial machinery of microhomology-mediated end joining (MMEJ) to human mtDNA editing, i.e., site-directed mutagenesis of the mitochondrial genome, a challenge that has not yet been fully resolved.

The 16.5 kb human mitochondrial genome encodes 13 transmembrane subunits of the respiratory chain complexes, 22 tRNAs, and two ribosomal RNAs required for the mitochondrial translation.

Therefore, any mutation in this extremely compact genome can lead to drastic changes in protein structure and/or expression, thereby causing respiratory chain dysfunction and decreased ATP production. Such defects particularly affect nervous and muscle tissue, but can also be associated with autoimmune diseases, cancer, diabetes, and many other disorders^41^. Despite substantial progress in understanding the molecular mechanisms of mitochondrial diseases, there is still a lack of treatments for patients affected by these disorders. The only notable exception is a recently approved in the European Union coenzyme Q analogue, idebenone, which has a positive effect in 50% early-stage LHON patients^42^.

Recent advances in direct editing of DNA sequences offer the possibility of converting one base pair into another, thus directly correcting certain pathogenic mutations^43^. The recently reported DdCBE-based approach of mammalian mitochondrial DNA editing utilized cytosine deamination and allowed efficient conversion of C:G into T:A pairs in tissues of living mice^10–12^. Similarly, conversion of A:T into G:C pairs has been achieved by applying mitochondria-targeted adenosine deaminase TALEDs^18,20,21^. Besides a significant level of off-target and bystander edits^13–15^. as well as context-dependence^10^, these editing approaches can introduce only transition nucleotide changes (purine-to-purine or pyrimidine-to-pyrimidine). The strategy we propose in the current study, which uses RMIS-based DNA oligonucleotide delivery and the intrinsic MMEJ enzymatic machinery for mammalian mitochondrial genome editing, is capable of introducing virtually any type of nucleotide changes including multiple mutations in the region of interest. For this, DNA oligonucleotides bearing regions of homology with mtDNA and containing desired mutations should be delivered into mitochondria of human cells.

### DNA import into mammalian mitochondria

Here, we demonstrated for the first time that RNA mitochondrial import signals (RMIS), previously discovered in our laboratory, can be used to target relatively long (∼ 50 bp) dsDNA molecules into human mitochondria. We are aware that the experimental estimation of DNA, as well as RNA, mitochondrial import can be biased^33^. For this reason, we treated isolated mitochondria and mitoplasts with a mixture of nucleases, optimized to remove possible DNA contaminants from the mitochondrial surface. We also performed control experiments by lysing a portion of mitochondria before nuclease treatment; the detection of DNA molecules in mitochondria and mitoplasts but not in the lysates confirmed their localization in the mitochondrial matrix (Figure 2). Interestingly, when we transfected cells with a DNA duplex carrying RMIS on a single strand, the entire duplex was detected in mitoplasts, indicating that the DNA strands were not separated during translocation across mitochondrial membranes (Figure 2D). This is consistent with our previous data, which showed that a tRNA could be imported into yeast mitochondria as a folded molecule^44^, and with previous studies demonstrating the import of dsDNA in plant and human mitochondria^26,27,45^.

The molecular mechanism of nucleic acid translocation across mitochondrial membranes is not fully understood. Polynucleotide phosphorylase (PNPase), located in the intermembrane space of mitochondria, may be involved in the recognition of stem-loop RNA structures (RMIS)^46–48^. Outer membrane translocase (TOM) and voltage-gated anion channel (VDAC) may facilitate translocation across the outer mitochondrial membrane (reviewed in^33^). It should be noted that VDAC oligomers have been reported to form pores for the release of mtDNA fragments^49^, and the structure and role of Porin1 hexamers have recently been studied in yeast^50^. We can thus hypothesize that pores formed by VDAC hexamers may serve not only for the export of mtDNA fragments, but also for the uptake of dsDNA fragments from the cytosol.

### Mitochondrial genome editing by donor DNA delivery

We demonstrated here that donor DNA delivery into human mitochondria results in a low but statistically significant introducing of designed nucleotide changes into intact mitochondrial loci (Figure 3). This donor DNA-induced editing without the preliminary cleavage of mtDNA can be explained in two ways: first, by occasional random mtDNA cleavage by reactive oxygen species (ROS)^51^ or by stalling of the replication fork^52^. These events, if occurred in the region of homology with donor DNA, can induce mtDNA reparation by MMEJ mechanism resulting in the designed nucleotide changes which have been introduced into the donor DNA oligonucleotides. The second possibility consists in the proposed MME model. The model of the MME is shown in Figure 5. We hypothesize that RNA hairpin structures can be removed from the donor DNA molecules by mitochondrial nucleases EndoG, which digests both DNA^53,54^ and RNA^54^ ds-substrates with preference for C:G tracks, and ExoG, which can cleave at RNA-DNA junctions^55^. Noteworthily, the RMIS sequence has a CG track in the ds-site (Figures 2A and 4A). Afterwards, like in MMEJ mechanism, PARP1 recruits MRE11, BRCA1 and CtlP. These proteins, performing the 5’-3’ ends recession and production of 3’ overhangs, have been identified within human mitochondria^22^. Strand invasion, microhomology alignment and extension by DNA synthesis then can be facilitated by RAD51 and POLQ, which have both nuclear and mitochondrial localizations^56–59^. Non-homologous tails are removed by FLAP endonuclease FEN1, followed by DNA synthesis and end-joining by DNA ligase III^52,57^. This model is supported by our data, which demonstrated that DNA duplexes were more efficient for human mitochondrial genome editing than the single-stranded chimeric oligonucleotides (Figure 3). This can be explained not only by the rapid degradation of single-stranded DNA, but also by the fact that the ends of donor double-stranded DNA molecules can mimic double-strand breaks in mtDNA, thereby recruiting PARP1 and the entire MMEJ machinery.

**Figure 5.**
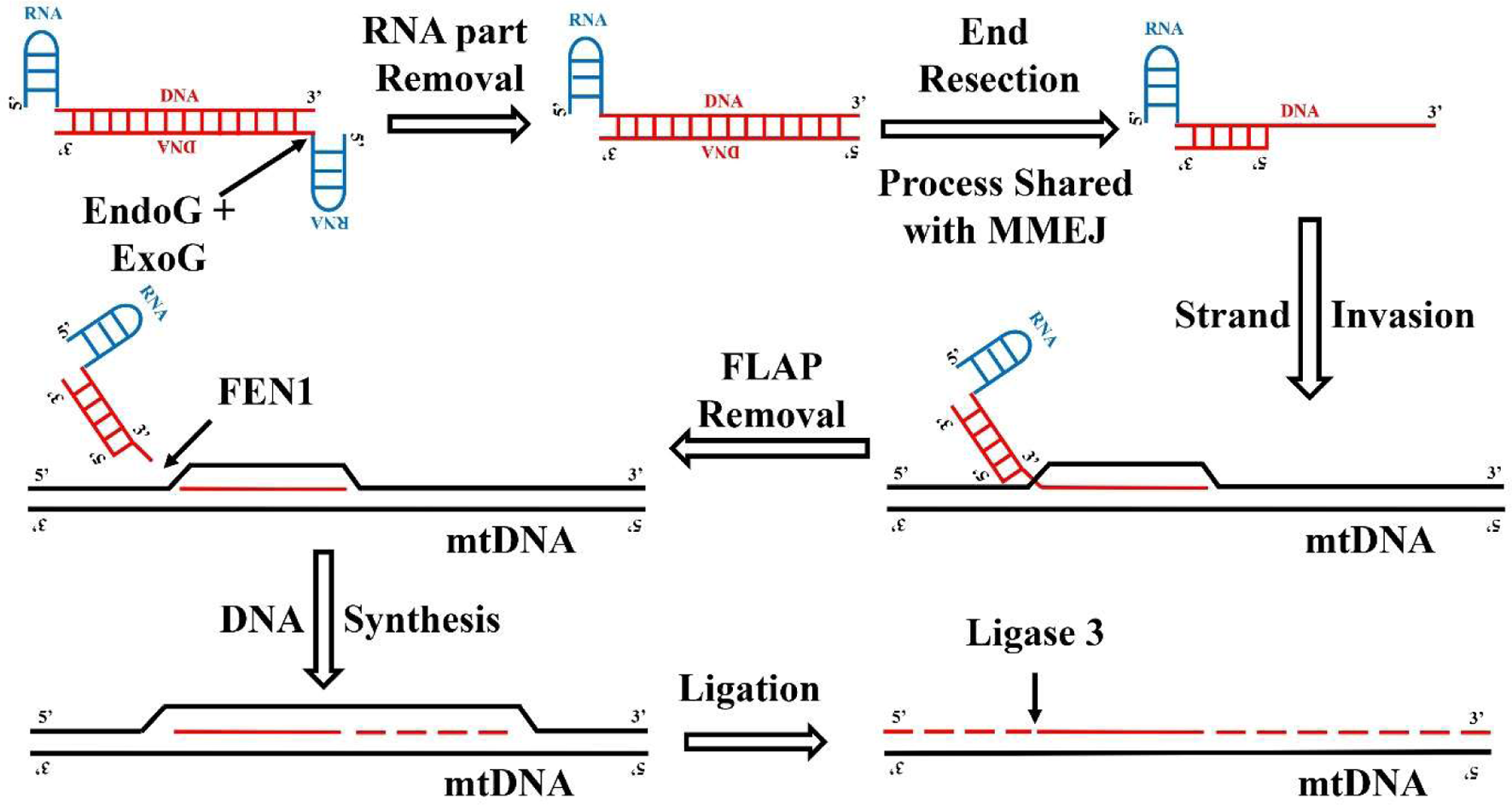
Hypothetic mechanism of microhomology-mediated mtDNA editing (MME). The process may start with RMIS removal (likely mediated by EndoG and ExoG nucleases), followed by end resection, which could be executed by mitochondrial MMEJ machinery, and 3’- overhangs formation. The next steps may involve strand invasion and FLAP removal by FEN1, followed by DNA synthesis and ligation by DNA ligase III. Newly synthesized DNA is represented by the red dashed line.

Both possible mechanisms are characterized by very low efficiency of introducing nucleotide changes (2×10^-4^–4×10^-4^ or less). Noteworthily, these values are close to the reported efficiency of introducing changes into the intact mammalian nuclear genome of somatic cells by homologous recombination (5×10^-4^)^60^. One possible way to improve the efficiency of our approach is to introduce site-specific cleavage into mitochondrial DNA. Double-strand breaks in editing mitochondrial sites are expected to facilitate integration of delivered oligonucleotides into these sites, enhancing efficiency of mammalian mitochondrial genome editing. Quick degradation of linearized unedited mitochondrial genomic DNA^23,24^ should further improve efficiency of our approach. Up to date, four types of DNA endonucleases have been successfully delivered into mitochondria and used for highly efficient site-specific mitochondrial DNA cleavage: (1) restriction enzymes (REs)^61–63^, (2) zinc-finger nucleases (ZFNs)^64–67^, (3) transcription activator-like effector nucleases (TALENs)^68–70^, and (4) the meganuclease ARCUS^71,72^. The primary purpose of all these techniques is a heteroplasmy shift, which is supposed to improve the ratio of wild-type to mutant mitochondrial genomic DNAs. Obviously, the heteroplasmy shift approach cannot be applied to nearly-homoplasmic mitochondrial diseases such as LHON. Nonetheless, a combination of our RMIS-based DNA oligonucleotide delivery approach with mitochondria-targeted site-specific endonucleases promises to be applicable for homoplasmic mitochondrial diseases as well. Importantly, this combined approach has the potential to introduce not only point changes into mammalian mitochondrial genomic DNA, but also large deletions and insertions.

Recently, the applications of CRISPR technology for specific mitochondrial genome cleavage have been reported^35,39^. We, therefore, attempted to combine CRISPR/mito-AsCas12a technology with donor DNA delivery to improve mitochondrial genome editing efficiency (Figure 4). A statistically significant number of the expected five-nucleotide change was detected in transfected cells; however, global editing was unexpectedly less efficient. This could be explained by lower respiration capacity of the cell line and/or also by a very low efficiency of mtDNA specific cleavage by the CRISPR/mito-AsCas12a system^39^. Notably, in these experiments, the engineered nucleotide changes were distributed over a larger region of the mitochondrial genome (16 nucleotides) (Figure 4A), which could also significantly reduce the editing efficiency. We believe that optimizing the mitochondrial CRISPR/mito-AsCas12a system together with donor DNA design can create an efficient mtDNA editing tool.

In summary, we report a novel strategy of mtDNA editing in living human cells via RMIS-based DNA oligonucleotide delivery. This approach was used for the predefined change of four or five nucleotides in the sequence of the human mitochondrial *ND4* gene. MME can introduce all possible changes into the mammalian mitochondrial genome, opening opportunities for mtDNA editing in basic and medical research.

## Materials and Methods

### Cell Culture

HEK293T cells (ATCC, Manassas, VA) were maintained in DMEM-high glucose medium (cat # 01-052-1A, Biological Industries) complemented with 4 mM L-Glutamine, 10% fetal bovine serum (FBS), and antibiotics (Penicillin G – 100 U/ml; Streptomycin Sulfate –100µg/ml; Nystatin – 12.5 U/ml). 1 mM pyruvate and 50 µg/ml uridine were added to the medium in order to culture HEK293T cells after mitochondrial genome editing.

Generation of the T-REx-293-Su9-AsCas12a cells with tetracycline-inducible expression of Su9-AsCas12a and culturing conditions for these cells are reported by us elsewhere^39^. When required, Su9-AsCas12a expression was induced by adding 100 ng/mL of tetracycline to culturing medium.

### Transfections

All transfection with HEK293T cells were conducted with the lipofectamine transfection reagent (cat # 18324012, ThermoFisher Scientific) according to the manufacturer instructions. Briefly, HEK293T cells were seeded in DMEM-high glucose medium complemented with 4 mM L-Glutamine, 10% fetal bovine serum (FBS) without any antibiotics a day before transfections. The HEK293T cells were transfected at confluence 80-90%. Transfection mixtures of the lipofectamine reagent with oligonucleotides were prepared according to the manufacturer instructions in DMEM-high glucose medium without serum and antibiotics. 30 µl of the lipofectamine reagent and 500 pmol of an ss-oligonucleotide or 250 pmol of an oligonucleotide duplex were used per one transfection in a T25 flask. 6 µl of the lipofectamine reagent and 100 pmol of an ss-oligonucleotide or 50 pmol of an oligonucleotide duplex were used per one transfection in a well of a 12 well plate. The medium on HEK293T cells was changed DMEM-high glucose medium without serum and antibiotics and the transfection mixtures were added to the cells. The medium on the transfected HEK293T cells was changed back DMEM-high glucose medium complemented with 4 mM L-Glutamine, 10% fetal bovine serum (FBS) without antibiotics in 4 hours after transfection. The next day after transfections the medium on the cells was changed to the same medium with antibiotics.

Transfections of T-REx-293-Su9-AsCas12a cells were conducted as described in the subsection “Mitochondrial Genome Editing Assay with Donor DNA Delivery and CRISPR/Cas12a”.

### Mitochondria Purification

Mitochondria purification for the MMEJ assay was conducted according a shorten version of a published protocol^73^ with modifications. Briefly, HEK293T cells were collected by trypsinization from 3 T75 flasks and washed with PBS. Then they were suspended in 2 ml of an ice-cold breakage buffer B_cells_-1 (225 mM mannitol, 75 mM sucrose, 0.1 mM EGTA, and 30 mM Tris-HCl pH 7.4) and disrupted by one out of two methods. The cells were homogenized either with 400 strokes at 4,000rpm applying Dounce-type homogenizer with Teflon pestle, or by passing the cells 25-30 times through a 25-gauge needle attached to a 2 ml syringe. The homogenate was centrifuged 3 times at 600g for 5 minutes at +4°C in order to remove nuclei, unbroken cells, and cell debris. The mitochondria containing supernatant was transferred to a new Eppendorf tube after each centrifugation. 1/20 of the nuclei-containing pellet after the first centrifugation was lysed in a mitochondrial lysis buffer (50 mM Tris-HCl pH 7.5, 100 mM NaCl, 10 mM MgCl_2_, 10% glycerol, 2 mM EGTA, 2 mM EDTA, 0.2% Triton X-100, and 1 mM DTT) complemented with Half Protease and Phosphatase Inhibitor Cocktail (cat # 78441, ThermoFisher Scientific) by suspending the pellet in 100µl of the ice-cold mitochondrial lysis buffer and leaving the suspension on ice for 30 minutes and centrifuging it at 13,200g for 5 minutes at +4°C. Protein concentration in the supernatant (nuclear extract) was assessed by the Bradford method and then it was aliquoted by 10 µl, snap frozen in liquid nitrogen, and kept at -80°C. Mitochondria was precipitated from the mitochondria-containing supernatant by centrifugation at 7,000g for 10 minutes at +4°C. The mitochondrial pellet was suspended in 2 ml of ice-cold buffer B_cells_-2 (225 mM mannitol, 75 mM sucrose, and 30 mM Tris-HCl pH 7.4) and centrifuged again at 7,000g for 10 minutes at +4°C. The mitochondrial pellet was again suspended in 2 ml of ice-cold buffer B_cells_-2 and centrifuged at 10,000g for 10 minutes at +4°C. The resulting mitochondrial pellet was suspended in 100µl of the ice-cold mitochondrial lysis buffer with Half Protease and Phosphatase Inhibitor Cocktail. The lysis was conducted and protein concentration was determined as it is described for the nuclear extract. The resulting mitochondrial extract was aliquoted by 10 µl, snap frozen in liquid nitrogen, and kept at -80°C. Mitochondria purification from HEK293T cells for the oligonucleotide delivery assay and deep sequencing was conducted as it was published elsewhere^32,36^. Briefly, HEK293T cells were detached and collected from one T75 flask by incubating them in 1 mM EDTA in PBS. Then these cells were washed in PBS and suspended in 1 ml of ice-cold buffer A (0.6 M sorbitol, 10 mM HEPES-KOH, pH 7.5, 1 mM EDTA) with 0.1% BSA. The cells were disrupted by passing them 30 times through a 25-gauge needle attached to an 1 ml syringe. The homogenate was diluted in 2 times by adding 1 ml of ice-cold buffer A with 0.1% BSA. The disrupted cells then were centrifuged 4 times at 600g for 5 minutes at +4°C in order to remove nuclei, unbroken cells, and cell debris. The mitochondria containing supernatant was transferred into a new Eppendorf tube each time. The mitochondria were precipitated at 15,000g for 20 minutes at +4°C. The mitochondrial pellet was then treated as described in subsections “Oligonucleotide Delivery Assay” and “Mitochondrial Genome Editing Assay”.

### Primer and Molecular Size Ladder Labeling and Radioactive PCR

20 pmol of P7 primer (Table S1) was labeled in 20 µl of a labeling reaction with 20 units of polynucleotide kinase T4 (PNK T4) (cat # M0201, NEB) and 13.3 pmol of ATP-[γ-^32^P] (6,000 Ci/mmol, cat # NEG035C001MC, PerkinElmer) in the PNK T4 buffer. 0.2 pmol of the radioactively labeled P7 primer were used per 10 µl of a PCR reaction along with 5 pmol of each – nonradioactive P5 and nonradioactive P7 primers (Table S1). The radioactive PCR was conducted with Taq polymerase at the final concentration 0.1 U/µl and at the final dNTP concentration 0.2 pmol/µl each in a commercial Taq polymerase buffer (cat # B9004S, NEB). The PCR conditions are further specified in the subsection “MMEJ Assay”.

Oligonucleotides MMEJ-Dir (99nt), MMEJ-Rev (91nt), MMJE-Linker-Dir (66nt), MMEJ-L-Dir (54nt), and MMEJ-L-Rev (46nt) (Table S1) were labeled at the same conditions as the P7 primer and used as a molecular size ladder in the MMEJ assay.

### MMEJ Assay

The MMEJ assay was conducted as it was described by researchers from Sathees C. Raghavan’s group^22,74^ with modifications. Briefly, duplexes MMEJ-Ovr-L, MMEJ-Ovr-R, MMEJ-L, MMEJ-Linker, and MMEJ-R were obtained by annealing of sense and antisense oligonucleotides: MMEJ-Ovr-L-Dir + MMEJ-Ovr-L-Rev, MMEJ-Ovr-R-Dir + MMEJ-Ovr-R-Rev, MMEJ-L-Dir + MMEJ-L-Rev, MMEJ-Linker-Dir + MMEJ-Linker-Rev, and MMEJ-R-Dir + MMEJ-R-Rev, correspondingly. The oligonucleotide sequences can be found in Table S2. Overhangs on each side of every duplex (Figure 1 A, B) are included in order to reduce PCR artifacts. The MMEJ assay buffer composition was 50 mM Tris-HCl pH 7.6, 20 mM MgCl_2_, 10% PEG-3350, 1 mM ATP, 1 mM DTT. All MMEJ reactions were conducted in the volume = 20 µl. 5 µg of mitochondrial or nuclear extract were added to one MMEJ reaction. In the MMEJ assays with MMEJ-Ovr-L and MMEJ-Ovr-R duplexes each duplex was added in the final concentration 4 nM. In the MMEJ assays with MMEJ-L, MMEJ-Linker, and MMEJ-R duplexes, MMEJ-L and MMEJ-R duplexes were added in final concentrations 4 nM each, while MMEJ-Linker duplex was added in the final concentration 8 nM. The MMEJ reactions were conducted for 4 hours at 37°C and stopped by heat inactivation at 65°C for 20 minutes. Each MMEJ reaction was diluted in 2 times with water and 2 µl of the diluted MMEJ reaction were taken for radioactive PCR with P5 and P7 primers in the volume of 10 µl. The radioactive PCR was conducted as it is described in the subsection “Primer Labeling and Radioactive PCR”. Precise PCR conditions for the MMEJ assays with MMEJ-Ovr-L and MMEJ-Ovr-R duplexes were: 94°C – 3min; cycle - 94°C – 30sec, 60°C – 1 min, and 68°C – 1 min (16 cycles); 68°C – 5 min. Precise PCR conditions for the MMEJ assays with MMEJ-L, MMEJ-Linker, and MMEJ-R duplexes were: 94°C – 3min; cycle - 94°C – 30sec, 60°C – 1 min, and 68°C – 1 min (26 cycles); 68°C – 5 min.

In the case of MMEJ assays with MMEJ-Ovr-L and MMEJ-Ovr-R duplexes, the recombination product is detectable by the radioactive PCR with P5 and P7 primers (Figure 1A and C, upper panel at the left). 5 µl of each radioactive PCR reaction were immediately loaded and resolved on a 10% denaturing polyacrylamide gel (PAAG) with 7M urea. The gels were dried and exposed to a phosphorimager screen, which was then visualized on Typhoon phosphorimager (FLA 7000).

In the case of MMEJ assays with MMEJ-L, MMEJ-Linker, and MMEJ-R duplexes, we asked whether the MMEJ activity can be used to join DNA ends through a linker containing an enzyme restriction site. Introduction of a novel DNA sequence (e.g. *BamHI* site) is an indication of MMEJ. Design of oligonucleotide duplexes in this case was different (Figure 1B and D, upper panel at the right). One duplex (MMEJ-L) has a 22nt microhomology arm, identical to the sequence located in the human mitochondrial genome to the left of the six-nucleotide sequence “GTC**<u>G</u>**CA” and designated “HS-1” (Figure 1B and D, upper panel at the right). The bold underlined G in this sequence is G11778 and is in fact the most commonly mutated nucleotide in LHON^6,7^. The other duplex has a 22nt microhomology arm, which is designated “HS-2”, identical to the sequence located in the human mitochondrial genome (to the right of the above six-nucleotide sequence in Figure 1B and D, upper panel at the right). These duplexes (MMEJ-L and MMEJ-R) cannot recombine with each other but only with the linker MMEJ-Linker. Therefore, the radioactive PCR with primers P5 and P7 can detect a product of two MMJE-mediated recombination events with the *BamHI* site in the middle (Figure 1B and D, upper panel at the right). In order to detect this product of double recombination, the radioactive PCR reactions were diluted in 2 times with water and 2 µl of each diluted radioactive PCR reaction were digested with 20 U of *BamHI* (cat # R0136S, NEB) in the *BamHI* buffer in 12 µl of the final volume for 6 hours at 37°C. The control undigested samples were incubated for the same time at the same conditions but without *BamHI*. Then the samples were loaded and resolved on a 10% denaturing PAAG with 7M urea. The gels were dried and exposed to a phosphorimager screen, which was visualized on Typhoon phosphorimager (FLA 7000).

### Oligonucleotide Delivery Assay

Oligonucleotides, which are listed in Table S5, were used in the oligonucleotide delivery and mitochondrial genome editing assays. RMIS-Dir and RMIS-Rev hybrid oligonucleotides contain at their 5’-end a known mitochondrial RNA import signal – modified D-arm of yeast 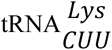 – rGrCrGrCrArArUrCrGrGrUrArGrCrGrC^29,30^. Duplex 1, duplex 2, duplex 3, and duplex 4 were obtained by annealing of oligonucleotides RMIS-Dir + RMIS-Rev, RMIS-Dir + Rev, Dir + RMIS-Rev, and Dir + Rev correspondingly. HEK293T cells were transfected with oligonucleotides and duplexes in T25 flasks as it is described in the subsection “Cell Culture and Transfections”. Immediately after transfections, the medium on the cells was changed to the medium with pyruvate and uridine. In 24 hours after transfections the transfected HEK293T cells were reseeded from T25 flasks to T75 flasks (one T25 flask to one T75 flask). In 72 hours after the transfections mitochondria were purified from the transfected cells as it is described in the subsection “Mitochondria Purification”. The purified mitochondria were divided in 6 equal parts and subjected to 10-minute incubation in 100 µl of one of the following buffers – the breakage buffer A (0.6 M sorbitol, 10 mM HEPES-KOH, pH 7.5, 1 mM EDTA), the swelling buffer B (10 mM HEPES-KOH, pH 7.5, 1 mM EDTA), and the lysing buffer C (10 mM HEPES-KOH, pH 7.5, 1 mM EDTA, 0.5% n-dodecyl-β-maltoside) with and without a mix of nucleases. Compositions of the buffers were already published elsewhere^32,36^. The mix of nucleases, which was added to each sample, consisted of 5 µl of the RNase A/T1 mix (2 mg/ml, 5,000 U/ml), 0.5 µl of RNase I_f_ (50,000 units/ml), 5 µl of DNase I (2,000 units/ml), 5 µl of Exo I (20,000 units/ml), and 0.25 µl of Exo III (100,000 units/ml). Immediately after the treatment, 1.07 ml of TRI reagent (cat # 93289, Sigma-Aldrich) with adjusted pH was added to each sample. The TRI reagent pH was adjusted by adding 70 µl of 3M KOH to each milliliter of the TRI reagent in order to ensure that the TRI reagent pH is higher than 8.0. This pH allows to purify the total nucleic acids (RNA + DNA). The total nucleic acids were then purified according to the manufacturer protocol for RNA purification. 60 µg of glycogen were added to the aqueous phase of each sample during the purification in order to ensure more complete oligonucleotide recovery. The purified total nucleic acids were subjected to the small RNA/DNA Northern blot as it is described in the subsection “Small RNA/DNA Northern Blot” in order to assess oligonucleotide delivery into mitochondria.

### Mitochondrial Genome Editing Assay with Donor DNA Delivery only

HEK293T cells in wells of a 12 well plate were transfected with oligonucleotides and duplexes, which were described in the subsection “Oligonucleotide Delivery Assay”, as it is described in the subsection “Cell Culture and Transfections”. Immediately after transfections, the medium on the cells was changed to the medium with pyruvate and uridine. In 24 hours after transfections, a half of the transfected HEK293T cells from each well of the 12 well plate was seeded in a 35 mm cell culture dish. In 82 hours after the transfections, the transfected HEK293T cells from each 35 mm dish were seeded in one T75 and one T25 flasks. In 144 hours after the transfections the transfected HEK293T cells in T25 flasks were frozen while mitochondria were purified from T75 flasks as it is described in the subsection “Mitochondria Purification”. The purified mitochondria were lysed in 200 µl a lysis buffer (10 mM Tris-HCl pH 8.8, 1 mM EDTA, 0.5% Triton X-100) by heating the samples at 95°C for 5 minutes. 2 µl of each lysate were used to amplify the edited site of the human mitochondrial genome with primers Mit-ND4-DS-Dir and Mit-ND4-DS-Rev (Table S1) applying Q5 high-fidelity DNA polymerase (Cat. # M0491, *New England Biolabs*, USA). The resulting PCR products were submitted for library preparation and deep sequencing at the Genomics Applications Laboratory (Core Research Facility, Faculty of Medicine, The Hebrew University of Jerusalem, Jerusalem).

### Mitochondrial Genome Editing Assay with Donor DNA Delivery and CRISPR/Cas12a

For *MGME1* knockout (when required), T-REx-293-Su9-AsCas12a cells were transfected in suspension with 40 nM DsiRNA targeting *MGME1* (hs.Ri.MGME1.13.8, IDT) (Table S5) using Lipofectamine RNAiMAX (Invitrogen) in Opti-MEM (Gibco, Fisher Scientific). After 6 hours, the transfection medium was replaced with complete EMEM. Three days later, transfection was repeated under standard adherent cell conditions.

After additional 48 hours, cells were transfected in suspension with crRNA (320 ng/ml) targeting the *ND4* gene of human mtDNA and donor DNA (RMIS-Rev-New or RMIS-Dir-New:RMIS-Rev-New at 2 µg/ml) using Lipofectamine 2000 (Invitrogen) in Opti-MEM. After 6 hours, the medium was replaced with EMEM supplemented with 100 ng/ml tetracycline to induce Su9-AsCas12a expression.

Seven days later, cells were detached, lysed, and total DNA was extracted using the QIAamp DNA Mini Kit (Qiagen). 100 ng of each purified mitochondrial DNA sample were used to amplify the edited site of the human mitochondrial genome with introduction of unique molecular identifiers (UMIs) according to a published protocol^40^ with some modifications. Briefly, reactions of UMIs introduction were conducted with the NGS-ND4-Dir primer (Table S2). Primer concentration was 0.02 μM. Total reaction volume was 10 μl. Q5 Hot Start High-Fidelity 2X Master Mix (Cat. # M0494, New England Biolabs) was used for this reaction. The reaction conditions were: 98°C – 2 min, 59°C – 15 min, 65°C – 15 min, 72°C – 7 min. The reactions were purified from primers with were purified with GeneRead Size Selection Kit (Cat. # 180514, QiaGen, Germany) according to the manufacturer’s instruction. Two rounds of purification were applied. The edited site with introduced UMIs was further amplified from purified DNA with primers Ext Primer and NGS-ND4-Rev applying the Q5 Hot Start High-Fidelity 2X Master Mix (Cat. # M0494, New England Biolabs). PCR conditions were: 95°C – 3min; cycle - 95°C – 30sec, 59°C – 40sec, and 68°C – 40sec (16 cycles); 68°C – 7 min.

The obtained amplicons were submitted for library preparation and deep sequencing at the Genomics Applications Laboratory (Core Research Facility, Faculty of Medicine, The Hebrew University of Jerusalem, Jerusalem).

### Small RNA/DNA Northern Blot

Total nucleic acid samples were resolved on denaturing 10% PAAG with 7 M urea. The transfer was conducted in x0.5 TBE at 1 mA/cm^2^ on zeta-probe membrane (cat # 1620159, BioRad). After UV-crosslinking the membranes were hybridized with the following probes: Dir and Rev (Table S5), Anti-tRNA-Thr (specific to mitochondrial Thr tRNA), and Anti-5-8S RNA (specific to cytosolic 5.8S rRNA) (Table S6).

### Deep Sequencing

The deep sequencing of PCR products was conducted at the Genomics Applications Laboratory (Core Research Facility, Faculty of Medicine, The Hebrew University of Jerusalem, Jerusalem). Libraries were prepared according to the 16S metagenomic sequencing library preparation protocol^75^. In brief, the PCR products were purified with AMPure XP beads and indexes, which are listed in Tables S7 and S8, were introduced by 8 cycles of index PCR. 150nt single end sequencing was conducted on the Illumina machine “NextSeq 500” (cat # SY-415-1001, Illumina) using the Nextseq 500/550 mid output kit v2.5 (150 cycles). The deep sequencing data are available in the Sequence Read Archive (SRA) database: https://www.ncbi.nlm.nih.gov/sra/PRJNA1332044.

The bioinformatic analyses of both sets of the deep sequencing data were conducted by Dr. Yuval Nevo (Info-CORE, Bioinformatics Unit of the I-CORE at the Hebrew University of Jerusalem, Jerusalem) as outlined below.

### Bioinformatic Analysis of Mitochondrial Genome Editing Assay with Donor DNA Delivery only

Briefly, sequence quality was inspected with FastQC software. Cutadapt software^76^ was employed in order to trim low-quality and adapter sequences. This software was further applied for filtering sequences shorter than 75nt and/or lacking the forward primer sequence. Remaining low-quality reads were removed with the fastq_quality_filter software. The refined reads were further aligned to the human genome version GRCh38 employing Bowtie 2 software^77^ with default parameters. Uniquely aligned reads with a single best alignment score, which span the expected amplicon mitochondrial genome positions (MT:11710-11855), were then inspected for changes in the editing site. Reads with the string “T0C0G1A” in their MD tag were designated as reads with edited sequence (“Mut”). Those of the remaining reads, which had an alignment score higher than -15, were designated as reads with wild type sequence (“WT”).

The Mut reads and eight the most similar human nuclear genomic sequences (Supplementary Table 6) were then aligned to the corresponding human mitochondrial genomic sequence applying the Bowtie 2 software with permissive parameters (version 2.3.4.3, command: bowtie2 -f -L 12 –local --mp 2 --rdg 3,1 --rfg 3,1 --all -x MT -U genomic_homologues.fa). The latter provided an unambiguous verification that most of the Mut reads are indeed better aligned to the human mitochondrial genomic sequence than to any human nuclear genomic sequence. This alignment also verified that most of the Mut reads have the predesigned change in the nucleotide sequence.

### Bioinformatic Analysis of Mitochondrial Genome Editing Assay with Donor DNA Delivery and CRISPR/Cas12a

Analogously to the previous analysis, the cutadapt software was applied to trim low quality, adapter, and poly-G sequences. The same software was utilized to remove reads, which became shorter than 95nt after trimming of low quality sequences. Fastq_quality_filter software was used for removal of remaining low-quality reads. The cleaned reads were aligned to the human genome version GRCh38 employing Bowtie 2 software with the -a parameter. Uniquely aligned reads with a single best alignment score, which span the expected amplicon mitochondrial genome positions (MT:11790-11803) in the expected orientation, were de-duplicated in a such way that each de-duplicated read has either a unique UMI or a unique human mitochondrial genome-aligned sequence. The unique aligned and de-duplicated reads were checked for the presence of any of the possible combinations of the 5 intended nucleotide substitutions. The unique aligned and de-duplicated reads with all the 5 intended nucleotide substitutions were considered as edited. The remaining reads were considered as wild type reads.

### Statistics

Statistical significance of differences in the mitochondrial genome editing efficiency was assessed by the two-sided permutation test with plus-one correction and adjustment via Benjamini-Hochberg false discovery rate. One simulation with 100,000 permutations was conducted for each pair of samples.

### Data availability statement

The deep sequencing data are available in the Sequence Read Archive (SRA) database: https://www.ncbi.nlm.nih.gov/sra/PRJNA1332044. The raw gel autography and Northern blot data are available from the corresponding authors.

## Acknowledgements

We thank Dr. Yuval Nevo from Info-CORE, the I-CORE Bioinformatics Unit of the Hebrew University of Jerusalem for Bioinformatics data analysis.

We thank Anne-Marie Heckel (UMR 7156 GMGM, Strasbourg) for technical help in cells manipulating.

N.E., N.S., N.N. and I.T. were supported by the Interdisciplinary Thematic Institute IMCBio+, as part of the ITI 2021-2028 program of the University of Strasbourg, CNRS and Inserm, supported by IdEx Unistra (ANR-10-IDEX-0002), EUR (IMCBio ANR-17-EUR-0016) and SFRI (STRAT’US project, ANR-20-SFRI-0012) within the framework of France 2030 National Program. N.N. was supported by Region Grand-Est (France). O.P. is supported by the German-Israeli Foundation for Scientific Research and Development (GIF; Grant No. 1561) and the German-Israeli Project Cooperation (DIP; Grant No. 17516).

