## Supplemental Tables 1-8 and Supplemental Figures 1 and 2 for "Mitochondrial genome microhomology-mediated editing by donor DNA delivery into mitochondria in human cells"

### Supplemental Material

**Supplemental Table 1**. Oligonucleotides Used in the MMEJ Assays. Annealing sites P5 and P7 primers are underlined. The *BamHI* site in oligonucleotides MMEJ-Linker-Dir and MMEJ-Linker-Rev is also underlined. Microhomology sites are in italic. All the other nucleotides are in bold.

| **Oligonucleotide Name** | **Oligonucleotide Sequence** |
|---|---|
| MMEJ-L-HS-Dir | **GACT**AATGATACGGCGACCACCGA<br>*CACTCACAGTCGCATCATAATC***ATGCATAA** |
| MMEJ-L-HS-Rev | **GCAT***GATTATGATGCGACTGTGAGTG*<br>TCGGTGGTCGCCGTATCATT |
| MMEJ-R-HS-Dir | **GATTAGAT***CACTCACAGTCGCATCATAATC*<br>TCGTATGCCGTCTTCTGCTTG**TGAG** |
| MMEJ-R-HS-Rev | CAAGCAGAAGACGGCATACGA<br>*GATTATGATGCGACTGTGAGTG***ATCT** |
| MMEJ-L-HS1-Dir | **GACT**AATGATACGGCGACCACCGA<br>*TCAAACTACGAACGCACTCACA***ATGCATAA** |
| MMEJ-L-HS1-Rev | **GCAT***TGTGAGTGCGTTCGTAGTTTGA*<br>TCGGTGGTCGCCGTATCATT |
| MMEJ-Linker-Dir | **CACAGTCC***TCAAACTACGAACGCACTCACA*<br>GGATCC*TCATAATCCTCTCTCAAGGACT***AGACGTGC** |
| MMEJ-Linker-Rev | **GTCT***AGTCCTTGAGAGAGGATTATGA* |

| | |
|---|---|
| | GGATCC*TGTGAGTGCGTTCGTAGTTTGA***GGAC** |
| MMEJ-R-HS2-Dir | **GATTAGAT***TCATAATCCTCTCTCAAGGACT*<br>TCGTATGCCGTCTTCTGCTTG**TGAG** |
| MMEJ-R-HS2-Rev | CAAGCAGAAGACGGCATACGA<br>*AGTCCTTGAGAGAGGATTATGA***ATCT** |
| MMEJ-Dir | GACTAATGATACGGCGACCACCGATCAAACTACGA<br>ACGCACTCACAGTCGCATCATAATCCTCTCTCAAG<br>GACTTCGTATGCCGTCTTCTGCTTGTGAG |
| MMEJ-Rev | CAAGCAGAAGACGGCATACGAAGTCCTTGAGAGAG<br>GATTATGATGCGACTGTGAGTGCGTTCGTAGTTTGA<br>TCGGTGGTCGCCGTATCATT |

**Supplemental Table 2**. Primers used in this study.

| Primer Name | Primer Sequence |
|---|---|
| P5 | AATGATACGGCGACCACCGA |
| P7 | CAAGCAGAAGACGGCATACGA |
| Mit-ND4-DS-Dir | TCGTCGGCAGCGTCAGATGTGTATAAGAGACAG<br>CCACGGGCTTACATCCTCAT |
| Mit-ND4-DS-Rev | GTCTCGTGGGCTCGGAGATGTGTATAAGAGACAG<br>GCGAGGCTTGCTAGAAGTCA |
| Ext Pimer | TCGTCGGCAGCGTCAGAT |
| NGS-ND4-Dir | TCGTCGGCAGCGTCAGATGTGTATAAGAGACAG |

| | |
|---|---|
| | NNNNNNNNNNNNGGGGTAAGGCGAGGTTAGC |
| NGS-ND4-Rev | GTCTCGTGGGCTCGGAGATGTGTATAAGAGACAG<br>CCCTCGTAGTAACAGCCATTC |

**Supplemental Table 3. Human Nuclear mitochondrial Sequences (NumtS) similar to the human mitochondrial sequence >chrM:11713-11853.**

| | |
|---|---|
| **Genomic Location 1** | **>chr5:134926943-134927083** |
| Genomic Sequence 1 | GTGAGGCTTGCTAGAAGTCATCAAAAGGCTATTAGTGGGA<br>GTAGGGTTTGAAGTCCTTGAGAGAGAATTATGATGCGACT<br>GTGGGTACGTTCGTAGTTTGAGTTTGCTAGGCAGAATAGTA<br>ATGAGGATGTAAGTCCGTGG |
| **Genomic Location 2** | **>chr5:100049269-100049409** |
| Genomic Sequence 2 | GCGAGGCTTGCCAGAAGTCATCAAAAGGCTATTAGTGGGAG<br>TAGGGTTTGAAGTCCTTGAGAGAGAATTATGATGCGGCTGT<br>GGGTTCGCTCGTAGTTTGAGTTTGCTAGGCAGAAAAATAAG<br>GAGGATGTAAGTCCGTGG |
| **Genomic Location 3** | **>chr7:64104212-64104352** |
| Genomic Sequence 3 | GTAAGATTTGCTATAAGTCTTCAAAAGGCTATTAGTGGGAG<br>TAGTGTTTGAAGCTCTCGAGAGAGTAATATGATTCGGCTAT<br>GGATTCGCTCATAGTTTGAATTTGCTAGGCAGAATAGTAAG<br>GACAAAGTAAGTCCATGG |
| **Genomic Location 4** | **>chr4:155455324-155455464** |

| | |
|---|---|
| Genomic Sequence 4 | GTAAGATTTGCTAGAAGTCATCAAGAGGCTATTAGCGGGA GCAGTGTTTGAAGGCCTCAGGTAAGAAATATGGTTTGGCTA TGGACTTGCTCGTAGTTTGAATTTGCTAGGCAGAAGAGTAA GGATGAAGTGAGTCCATGA |
| **Genomic Location 5** | **>chr8:67585876-67586015** |
| Genomic Sequence 5 | GTCTGATTTGCTAGAAGTCATCCTGAGGCTATTAGTGGGAGT AGTGTTTGAAGGCCTCAGTAAGTAATATGGTTCATCTATGGG CTCACTCATAGTTTGAGTTTGCTAGGCAGAATAGTAGGGAT GAAGTGAGTCCATAA |
| **Genomic Location 6** | **>chr4:25718362-25718490** |
| Genomic Sequence 6 | GTAAGATTTGCTAGAAGAGGCTATTAGTGGGAGCAGTGTTT GAAGCCCTCGGGTAAGTAATATGGTTCATCTATGGACTCGC TCATAGTTTGAATTTGCTAGGCAGAATAGTAAGGATGAAGT GAATGC |
| **Genomic Location 7** | **>chr16:10721979-10722119** |
| Genomic Sequence 7 | GTAAGATTTGCTAGAAGTCATCAGGAGGCTATTAGTGGAAG CAATGTTTGAAGGCCTTGGAGAAGTAGTATAATTCGGCTAT GGACTCGTTCATAGTTCCAATTTGCTAGGCAGAATAGTAAG GATGAAGTGAGTCCATGG |
| **Genomic Location 8** | **>chr1:235540322-235540462** |
| Genomic Sequence 8 | GTAAGATTTGCTAGAAGTCATCAAGAGGCTATTAGCAGAAG CAGTGTTTGAAGGCGTTCGGTAAGTAATAGGGTTCACCTAT GGGCCTGCTCGTAGTTTGAATTTGCTAGGCAGAATAGTAAG |

| | |
|---|---|
| | GCTGAAGTGAGTCCATGA |

**Supplemental Table 4. Human Nuclear mitochondrial Sequences (NumtS) similar to the human mitochondrial sequence >chrM:11643-11870.**

| **Genomic Location 1** | **>chr5:134926926-134927153** |
|---|---|
| Genomic Sequence 1 | CCCTCGTAGTAACAGCCATCCTCATCCAAACCCCCTGAAGC TTCACCGGCGCAGTCATTCTCATAGTCGCCCACGGACTTAC ATCCTCATTACTATTCTGCCTAGCAAACTCAAACTACGAAC GTACCCACAGTCGCATCATAATTCTCTCTCAAGGACTTCAAA CCCTACTCCCACTAATAGCCTTTTGATGACTTCTAGCAAGCC TCACTAATCTCGCCTTACCCC |
| **Genomic Location 2** | **>chr5:100049260-100049479** |
| Genomic Sequence 2 | CCCTCATAGTAACAGCTATTCTCATCCAAACCCCCTGAAGCT TCACCGGCGCAGTCATTCTCATAATTGCCCACGGACTTACAT CCTCCTTATTTTTCTGCCTAGCAAACTCAAACTACGAGCGAA CCCACAGCCGCATCATAATTCTCTCTCAAGGACTTCAAACCC TACTCCCACTAATAGCCTTTTGATGACTTCTGGCAAGCCTCG CCAACCTCGC |
| **Genomic Location 3** | **>chr7:64104195-64104401** |
| Genomic Sequence 3 | TCATTCAAACCCCCTGAAGCTTTACTGGTGCAATTACCCTCA TAATTGCCCATGGACTTACTTTGTCCTTACTATTCTGCCTAG CAAATTCAAACTATGAGCGAATCCATAGCCGAATCATATTA |

| | |
|---|---|
| | CTCTCTCGAGAGCTTCAAACACTACTCCCACTAATAGCCTTT<br>TGAAGACTTATAGCAAATCTTACTAACCTTGCCTTACCCC |
| **Genomic Location 4** | **>chr10:36434949-36435150** |
| Genomic Sequence 4 | CAAACCCCCTGAAGCTTTACTGGTGCAATTACCCTCATAATT<br>GCCCACAGACTTACTCTGTCCTTACTATTCTGCTTAGCAAAT<br>TCAAATTACGAGTGAGTCCACAGCCGAATTATACTGCTCTCT<br>TAACGCCTTCAACCATTACTTCCACTAATAGCTTTTTGATGA<br>CTTACAGCAAATCTTACCAACCTTGCCTTACCCC |

**Supplemental Table 5**. **Oligonucleotides, which were used in the oligonucleotide delivery and mitochondrial genome editing assays.** Uppercase letters, which represent ribonucleotide residues, are always preceded by a lowercase letter “r”. All the other uppercase letters represent deoxyribonucleotide residues. Ribonucleotide residues in chimeric RNA/DNA oligonucleotides are also highlighted in blue. Deoxyribonucleotide residues, which compose microhomology arms, are in italic. Deoxyribonucleotide residues, which differ from the human mitochondrial genome sequence, are in bold and underlined. Ribonucleotide residues of the crRNA, which are not complementary to the targeted site, are in bold.

| **Oligonucleotide Name** | **Oligonucleotide Sequence** |
|---|---|
| RMIS-Dir | rGrCrGrCrArArUrCrGrGrUrArGrCrGrC*CAAACTACGAA*<br>*CGCACTCACAG*__GAT__C__C__*TCATAATCCTCTCTCAAGGACT* |
| RMIS-Rev | rGrCrGrCrArArUrCrGrGrUrArGrCrGrC*AGTCCTTGAGA* |

| | |
|---|---|
| | *GAGGATTATGA***G**G**ATC***CTGTGAGTGCGTTCGTAGTTTG* |
| Dir | *CAAACTACGAACGCACTCACAG***GAT**C**C***TCATAATCCT CTCTCAAGGACT* |
| Rev | *AGTCCTTGAGAGAGGATTATGA***G**G**ATC***CTGTGAGTGC GTTCGTAGTTTG* |
| RMIS-Dir-New | rGrCrGrCrArArUrCrGrGrUrArGrCrGrC*CACTCACAGTCG CATCATAAT***T**CT**A**TC**A**CAAGG**C**CT**C***CAAACTCTACTCCC* |
| RMIS-Rev-New | rGrCrGrCrArArUrCrGrGrUrArGrCrGrC*GGGAGTAGAGTT TG***G**AG**G**CCTTG**T**GA**T**AG**A***ATTATGATGCGACTGTGAGTG* |
| crRNA | ***rUrArArUrUrUrCrUrArCrUrCrUrUrGrUrArGrArU***rArArG rUrCrCrUrUrGrArGrArGrArGrGrArUrUrA |
| siRNA MGME1 Direct Strand | rGrGrGrUrGrArArArGrUrArUrGrCrUrUrUrCrCrArArGrGrCrU rUrC |
| siRNA MGME1 Reverse Strand | rArGrCrCrUrUrGrGrArArArGrCrArUrArCrUrUrUrCrArCrCrC |

**Supplemental Table 6. Probes for small RNA/DNA Northern blot hybridization.**

| Probe Name | Probe Sequence |
|---|---|
| Anti-tRNA-Thr | TCTCCGGTTTACAAGAC |
| Anti-5-8S RNA | GGCCGCAAGTGCGTTCGAAG |

**Supplemental Table 7**. Description of samples for deep sequencing from the mitochondrial genome editing experiment with donor DNA only.

| Sample final name | Biological description | Index1 | Index2 |
|---|---|---|---|
| Sample_1 | Mock Transfection | AAGAGGCA | TCGCATAA |
| Sample_2 | Transfection 1 | GCTCATGA | ATAGCCTT |
| Sample_3 | Transfection 2 | ACTCGCTA | TCTTACGC |
| Sample_4 | Transfection 3 | GCGTAGTA | AGCTAGAA |
| Sample_5 | Transfection 4 | TACGCTGC | CGGAGAGA |

**Supplemental Table 8**. Description of samples for deep sequencing from the mitochondrial genome editing experiment with donor DNA and CRISPR.

| Sample final short name | Biological condition | Index 1 | Index 2 |
|---|---|---|---|
| Sample_CRISPR_1-1 | CRISPR + MGME1 siRNA | ACCAATTC | ACTAGAGT |
| Sample_CRISPR_1-2 | CRISPR + MGME1 siRNA | GGTGGACG | CATGAATG |
| Sample_CRISPR_1-3 | CRISPR + MGME1 siRNA | TGCGTCTC | GTTCATAG |
| Sample_CRISPR_2-1 | ssDNA + CRISPR | GGATTGTG | GTGGATAT |

| | | | |
|---|---|---|---|
| Sample_CRISPR_2-2 | ssDNA + CRISPR | GTTGCCGG | TTAAACCG |
| Sample_CRISPR_2-3 | ssDNA + CRISPR | CGTAAAGT | GGCGTTCA |
| Sample_CRISPR_3-1 | ssDNA + CRISPR + MGME1 siRNA | GGTGTGAA | GTACACTA |
| Sample_CRISPR_3-2 | ssDNA + CRISPR + MGME1 siRNA | AGGTTATC | TTCTAATG |
| Sample_CRISPR_3-3 | ssDNA + CRISPR + MGME1 siRNA | TCGATTGT | TTCCGTCA |
| Sample_CRISPR_4-1 | dsDNA | CTAGATAA | GAGCCTTT |
| Sample_CRISPR_4-2 | dsDNA | GGGTTCCG | CTCGAATC |
| Sample_CRISPR_4-3 | dsDNA | AAGTAAGA | AGAGGTCA |
| Sample_CRISPR_5-1 | dsDNA + CRISPR | TCACCCAT | TTCCCAAG |
| Sample_CRISPR_5-2 | dsDNA + CRISPR | AATATTCG | GCTTCCAG |
| Sample_CRISPR_5-3 | dsDNA + CRISPR | GTCCCGGA | GCATTCAC |
| Sample_CRISPR_6-1 | dsDNA + CRISPR + MGME1 siRNA | GTGCGGTG | TGACAGTC |
| Sample_CRISPR_6-2 | dsDNA + CRISPR + MGME1 siRNA | CCTTGACG | GTGACCTA |
| Sample_CRISPR_6-3 | dsDNA + CRISPR + MGME1 siRNA | CTTGACGA | TGAGTTGC |

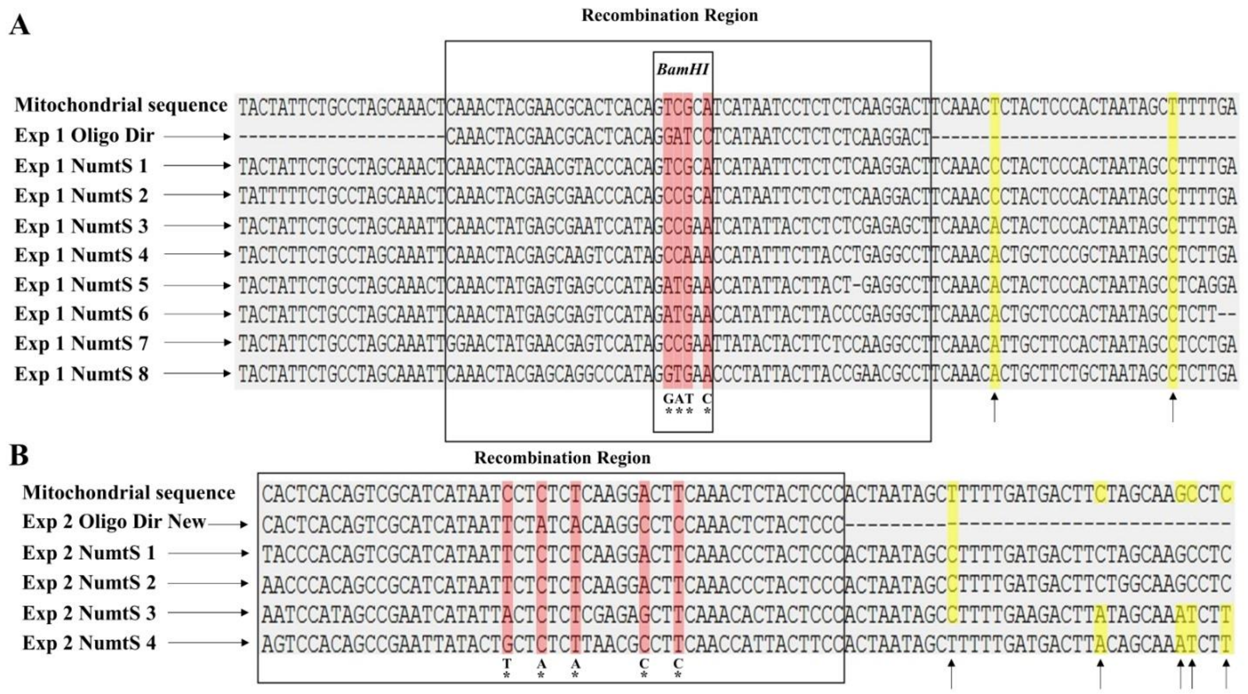


**Supplemental Figure 1. Alignment of the amplified human mitochondrial sequences to similar nuclear mitochondrial sequences (NumtS) at the sequenced amplicon region encompassing the edited *ND4* site.** Alignments of a part the amplified and sequenced human mitochondrial genomic region, harboring the edited *ND4* site, to the donor DNA sequences (Dir and Dir New) with the expected nucleotide edits and to similar human NumtS are shown. The alignments were conducted by the MAFFT software (https://mafft.cbrc.jp/alignment/server/index.html)[1,2]. Primers annealing sites for the amplified mitochondrial sequence and corresponding nuclear sequences are not included in the figure. The expected nucleotide edits are indicated at the bottom in bold and marked with asterisks. Corresponding nucleotides in the aligned sequences are highlighted in pink. The human nuclear sequences with the primers annealing sites can be found in Supplemental Tables 5 and 6. The recombination regions are surrounded by large rectangles. Nucleotide positions outside the recombination regions, at which the mitochondrial sequences differ from the corresponding nuclear sequences, are highlighted in yellow and marked by arrows at the bottom. (A) Alignment of the entire amplified and sequenced human mitochondrial genomic sequence without primers annealing sites from the experiment presented on Figure 3 to corresponding donor DNA and human NumtS is shown. The small rectangle surrounds sequence, which corresponds to the BamHI site in the edited mitochondrial DNA. (B) Alignment of the right part of the amplified and sequenced human mitochondrial genomic sequence without the primer annealing site from the experiment presented on Figure 4 to corresponding donor DNA and human NumtS is shown.

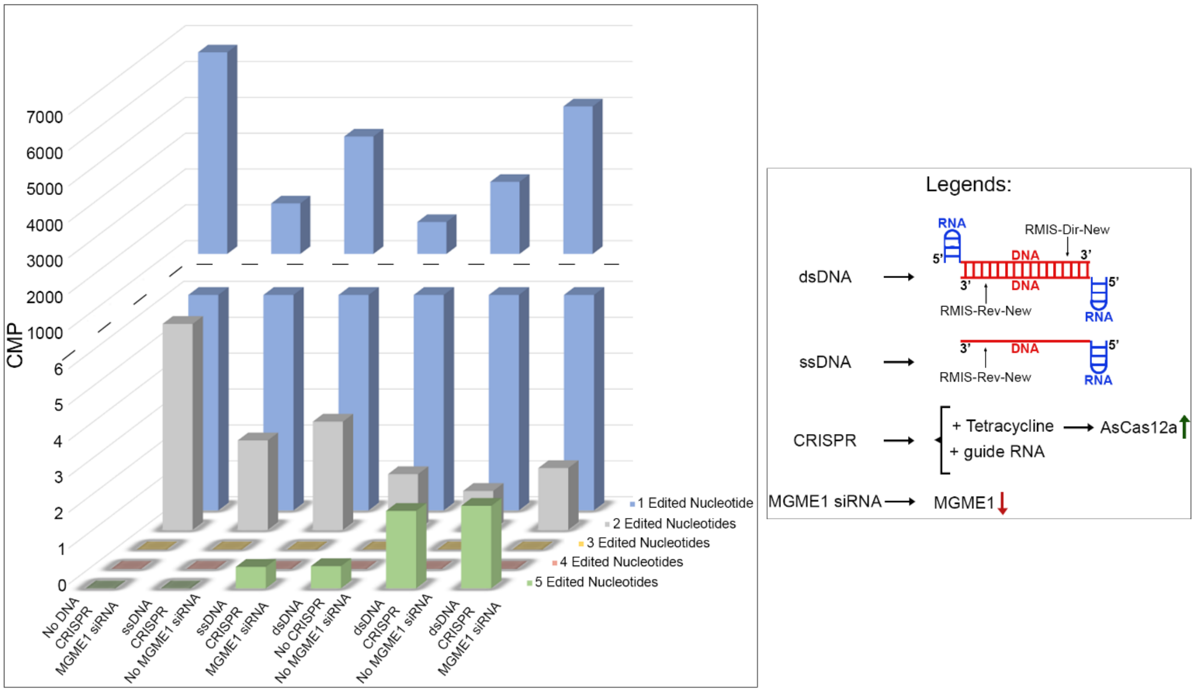


**Supplemental Figure 2. Introduction of changes in mtDNA by donor DNA delivery and CRISPR in mitochondria of T-REx-293-Su9-AsCas12a cells – supplemental information.** Left panel: counts of reads with five-, four-, three-, two-, or one-edited nucleotides per one million

(Counts Per Million = CPM) of reads unambiguously aligned to the mitochondrial sequence (MT:11643-11870) for each transfection are shown on the 3D graph. Right panel: legends, explaining transfections and treatments.

**Supplemental References.**


1. Katoh, K., Rozewicki, J. & Yamada, K. D. MAFFT online service: multiple sequence alignment, interactive sequence choice and visualization. *Brief Bioinform* **20**, 1160–1166 (2019).

2. Kuraku, S., Zmasek, C. M., Nishimura, O. & Katoh, K. aLeaves facilitates on-demand exploration of metazoan gene family trees on MAFFT sequence alignment server with enhanced interactivity. *Nucleic Acids Res* **41**, W22–W28 (2013).
